# Polycyclic aromatic hydrocarbons, gut microbiome composition, impulsivity, and attention covary in a human cohort

**DOI:** 10.64898/2026.03.02.709058

**Authors:** Austin J. Hammer, Kristin Kasschau, Ed Davis, Peter Hoffman, Katie Johnson-Camacho, Alexandra Alexiev, Kim Anderson, Jackilen Shannon, Lisa K. Marriott, Thomas J. Sharpton

**Affiliations:** Department of Microbiology, Oregon State University, Corvallis, OR, USA; Center for Quantitative Life Sciences, Oregon State University, Corvallis, OR, USA; Department of Environmental and Molecular Toxicology, Oregon State University, Corvallis, OR, USA; Department of Medicine, University of California Los Angeles, Los Angeles, CA, USA; Knight Cancer Institute, Portland, OR, USA; School of Medicine, Oregon Health & Science University, Portland, OR, USA; School of Public Health, Oregon Health & Science University, Portland, OR, USA; Department of Statistics, Oregon State University, Corvallis, OR, USA; Linus Pauling Institute, Oregon State University, Corvallis, OR, USA

## Abstract

Polycyclic aromatic hydrocarbons (PAHs) are pervasive environmental pollutants linked to adverse neurobehavioral outcomes, yet the biological pathways coupling exposure to behavior are poorly defined. The gut microbiome is both sensitive to PAH exposure and a modulator of central nervous system function, suggesting it may mediate how PAH exposure influences neurobehavior. We tested whether PAH exposure, gut microbiome composition, and neurobehavioral function covary in a statewide sample of 34 adults stratified into high-impulsivity/poor-attention (HH) and low-impulsivity/fast-attention (LL) groups. Participants provided fecal samples for 16S rRNA profiling and wore silicone wristbands for 30 days to passively sample PAH exposure. Higher PAH exposure associated with HH group membership in a sex-dependent manner, with the largest elevations among HH males. At the community level, PAH exposure profiles correlated with microbiome dissimilarity, and HH membership associated with increased alpha-diversity and altered community composition relative to LL members. At the taxon level, 21 genera were significantly associated with 14 PAH compounds (FDR < 0.1). No individual genera were significantly associated with neurobehavioral group after multiple testing correction. Nevertheless, cross-referencing PAH-responsive genera (FDR < 0.1) against those with nominal neurobehavioral associations (p < 0.05) identified two candidate genera, *Hydrogenoanaerobacterium* and *Methanobrevibacter*, whose abundance covaries with both PAH exposure and neurobehavioral phenotype. Both have been independently linked to cognitive or neurological outcomes in prior work. These findings support a three-way relationship among environmental chemical exposure, gut microbiome composition, and neurobehavioral function, establishing an empirical foundation for testing microbiome-mediated links between PAH exposure and neurobehavioral outcomes.

**IMPORTANCE:** PAH exposure is widespread and associates with impulsivity and attention problems, but how exposure translates into neurobehavioral risk is unclear. The gut microbiome is a plausible intermediary: gut microbes biotransform environmental chemicals and produce metabolites that influence brain function. In a statewide adult cohort, we show that higher PAH exposure tracks with greater impulsivity and poorer attention in a sex-dependent manner, and that both PAH exposure and neurobehavioral phenotype associate with distinct gut microbiome features at the community and taxon levels. We identify candidate genera at the intersection of PAH exposure and neurobehavioral group whose biology independently implicates them in cognitive and neurological function. By demonstrating that all three domains covary within a single human cohort, this work moves beyond pairwise associations to identify candidate microbial intermediaries for mechanistic investigation. Defining the microbiome constituents that respond to PAH exposure and co-associate with neurobehavioral phenotypes creates opportunities to test microbiome-targeted or exposure-reduction strategies for mitigating PAH-related neurobehavioral impacts.

## INTRODUCTION

Polycyclic aromatic hydrocarbons (PAHs) are a group of persistent and ubiquitous environmental pollutants that are principally produced during incomplete combustion of organic matter, such as during the burning of fossil fuels and biomass (1–3). These persistent organic pollutants can linger in a variety of environmental locales, such as air, water, soil, and food, and are known to bioaccumulate in organisms (4–7). PAHs readily cross the blood-brain barrier due to their lipophilic nature (8) and exert especially toxic effects on the brain and neural tissue, where they are linked to increased susceptibility to cognitive and neurological disorders (9–11). Research investigating prenatal and early postnatal exposure to PAHs has consistently reported negative impacts on childhood behavioral development, including hyperactivity and attention deficits (12–15). Higher levels of PAH exposure in children correlate with increased inattentiveness and hyperactivity (12–14), and exposure to PAHs and other xenobiotics correlates with impulsive behavior in both children and adults (13, 16, 17). Notably, PAH metabolism shows pronounced sex-dependent variation (18, 19), suggesting that the neurobehavioral consequences of exposure may differ between males and females. Despite clear links between PAH exposure and neurobehavioral outcomes, the intermediate biological systems that connect exposure to behavioral change remain poorly characterized, particularly in humans (20, 21).

The gut microbiome is emerging as a plausible intermediary in this pathway. Recent research demonstrates that the gut microbiome is sensitive to PAH exposure and may play a critical role in mediating the toxicity of xenobiotic compounds including PAHs (22, 23). Gut microorganisms possess the enzymatic capacity to metabolize PAHs, and this metabolism can produce both toxic (24) and non-toxic byproducts (25, 26) that exert their own effects on host physiology (27), including bioactive compounds such as estrogenic metabolites (28, 29). Many gut taxa display altered abundance following broad PAH exposure (30, 31), and studies in animal models demonstrate that exposure to the PAH benzo[a]pyrene (BaP) shifts the overall composition of the gut microbiota (32–35). Critically, in zebrafish, the microbiome has been shown to mediate the impact of BaP on neurodevelopment (35), providing direct evidence that gut microbes can modulate PAH-driven neurobehavioral outcomes. Independent of PAH exposure, gut microbiome composition is linked to behavioral variation in human clinical studies (36–38), and gut microbes modulate production of neurologically relevant metabolites, including short-chain fatty acids and neurotransmitters such as serotonin, that influence neural function (39–41). Taken together, these lines of evidence support the hypothesis that the gut microbiome is a biological intermediary between environmental PAH exposure and neurobehavioral outcomes, through its capacity to biotransform PAH compounds into bioactive metabolites, its sensitivity to PAH-induced compositional disruption, and its established influence on brain function and behavior.

Prior work has established pairwise links between PAH exposure and the gut microbiome, PAH exposure and impulsivity/attention, and the microbiome and behavior, but these three factors have not been examined simultaneously within a single cohort. Our study addresses this gap by testing whether PAH exposure, gut microbiome composition, impulsivity, and attention converge in humans. Adults were recruited from a large statewide population-based cohort study (42, 43) into the Exposure, Microbiome, and Cognition Cohort (EMC2), stratified into groups with concordantly high impulsivity and poor attention (HH) or concordantly low impulsivity and fast attention (LL). We integrated passively sampled PAH exposure profiles from silicone wristbands, 16S rRNA gut microbiome profiles, and neurobehavioral assessments to jointly examine the relationship between PAH exposure, gut microbiome composition, and neurobehavior in 34 participants with complete paired data. Overall, our analyses find that higher PAH exposure associates with HH group membership in a sex-dependent manner, that both PAH exposure and neurobehavioral group track with shifts in gut microbiome composition, and that specific microbial genera respond to PAH exposure at the taxon level. Together, these results provide evidence that environmental chemical exposure, the gut microbiome, and neurobehavioral function form an interacting system within a single human cohort.

## MATERIALS AND METHODS

### Study Context and Setting

This multi-site cross-sectional study engaged adults in an *Exposure, Microbiome and Cognition Cohort* (EMC2; OHSU IRB study #23221; eCRIS #6688). The EMC2 study leveraged established research infrastructure from the Healthy Oregon Project (HOP; OHSU IRB #18473), which afforded access to a large, geographically distributed population who was already familiar with and adherent to clinical study practices. The Healthy Oregon Project (www.healthyoregonproject.org) is an Oregon-wide population-based cohort that supports basic and clinical research, and importantly, aims to influence the health of Oregonians through providing no-cost genetic screening for heritable cancers (42). HOP employs a cost-effective remote recruitment approach that establishes a large-scale cohort for population-based studies (43).

#### EMC2 Study Design for Participant Recruitment

The EMC2 pilot study leveraged HOP’s research infrastructure for participant recruitment, selection, and data collection in compliance with both IRB-approved protocols. Due to budgetary limits, a sample of 50 individuals was recruited for this pilot study by email, 34 of whom provided complete paired data across all three domains (fecal microbiome, wristband PAH exposure, and neurobehavioral assessment; Table S1). Targeted study recruitment for cases and controls leveraged neurobehavioral data to identify highest (HH) and lowest (LL) quartiles of impulsivity and attention. Participants in HH or LL recruitment groups were emailed by the HOP study team about the study opportunity. EMC2 participants consented via HOP app to provide a fecal sample for microbiome analysis and submit their worn silicone-based wristbands to quantify participants’ passively accumulated PAHs from the environment. This study’s eligibility criteria excluded individuals with antibiotics use in the past six months. Inclusion criteria were 18 years of age or older, consent to the HOP repository, creation of a secure Medable account, completion of study behavioral assessments, and consent to this study.

#### Behavioral Assessments

All HOP participants are invited to complete data collection modules that assess lifestyle behaviors and health risks (43), including neurobehavioral measures of impulsivity and attention used to select participants for the EMC2 study. Quartiles for each measure identified potential participants for case-control groups.

##### Impulsivity

HOP participants completed the Barratt Impulsiveness Scale (BIS-15) (44), a fifteen-item instrument scored on a 4-point Likert scale to assess total impulsivity in clinical and non-clinical populations (45). Published work from our team details BIS-15 implementation with adult (46) and adolescent (47) populations. BIS-15 includes attentional, motor, and non-planning facets of impulsivity, with total scores ranging 15–60. Quartiles applied previously defined cut points for Q1 (15–28, least impulsive), Q2 (29–33), Q3 (34–37), and Q4 (38–60, most impulsive) to data from 3,106 HOP participants who completed BIS-15 at the time of EMC2 recruitment (Table S2).

##### Attention

Participants completed a validated 3-minute version of the Psychomotor Vigilance Task (PVT) (48), termed PVT-B (49). PVT-B provides an objective measure of reaction time (RT), errors of omission (i.e., lapses, lack of response in sufficient time), and errors of commission (i.e., false starts, where participants provide a response without a stimulus). At the time of EMC2 recruitment, 2,678 participants completed PVT-B, with individuals’ median reaction time used to assign quartile cut points. Individuals with RTs >1,300 msec were excluded as outliers (n=3; 2–16x higher than remaining data), with quartiles comprising Q1 (≤14 msec, lowest and fastest reaction times), Q2 (315–339 msec), Q3 (340–370 msec), and Q4 (≥371 msec, longest and slowest reaction times; Table S2).

##### Defining case-control groups for recruitment

Cases were defined as participants in the highest quartiles (Q4) for both impulsivity and attention, termed high/high (HH) to refer to individuals with high impulsivity and long reaction times. Controls in this study represented lowest quartiles (Q1), designated as low/low (LL) to refer to participants with low impulsivity and low (i.e., fast) reaction times. A total of 2,628 participants completed both instruments, with quartile distributions shown in Table S2, with no difference between group sizes (Chi-Square p=0.33). Once removing participants who were missing sex, age, impulsivity data, or reaction times, a total of 422 participants qualified for LL or HH group recruitment, with ages comparable between groups (p=0.77; LL=40.4 years ±11.8 SD; HH=40.8 ±12.0). Controls (LL=283, 79.5% female) and cases (HH=139, 85.6% female) had similar sex distributions (p=0.12). The relationship between impulsivity and attention in the EMC2 recruitment sample was confirmed by independent samples t-tests, with significant differences observed between impulsivity quartiles (i.e., Q1 vs Q4) for all attention metrics studied (p<0.001, Table S3). In addition to higher reaction times, participants in the most impulsive quartile (Q4) had more lapses and fewer false starts, consistent with less attention and slow response times. Thus, study controls represented lowest quartiles (LL) of impulsivity and response latency (i.e., more attention), while the highest quartiles (HH) represent participants with high impulsivity and higher (i.e., slower) reaction times indicative of worse attention.

### Passive PAH Exposure Sampling using Silicone Wristbands

#### Preparation and deployment

To measure each participant’s PAH exposure, we prepared silicone wristbands (24hourwristbands.com, Houston, TX, U.S.) that enable passive sampling of 34 distinct PAHs (50). We rinsed wristbands with deionized water and used a vacuum oven (300°C for 12 hours at 0.1 Torr; Blue M vacuum oven, no. POM18VC; Welch® DuoSeal pump, no. 1405, Mt. Prospect, IL, U.S.) to remove manufacturing process-related chemicals in the silicone polymer. Clean wristbands were mailed to participants, who wore the wristbands for 30 consecutive days prior to stool sampling (below). Used wristbands were then sealed and shipped back to us. We collected and analyzed quality control (QC) samples from each group of prepared wristbands to ensure all wristbands met our data quality objectives (51).

#### Cleaning and extraction

We cleaned wristbands that met our inclusion criteria and extracted the chemicals from the silicone as previously described (52). Particles on the wristband surface do not reflect the chemical fraction available to be absorbed by the body, and we removed particles with two rinses of 18 MΩ·cm water and one rinse of isopropanol (50, 52). We extracted chemicals by adding two separate 50 mL volumes of ethyl acetate at room temperature to the wristbands. We quantitatively concentrated the ethyl acetate to one mL using TurboVap® 500 closed cell evaporators and a TurboVap® LV evaporator workstation (Biotage LLC, Charlotte, NC, U.S.).

#### Chemical analysis

We quantitatively analyzed all wristband extracts for 64 PAHs and alkylated PAHs. Analysis was performed using an Agilent 7890B gas chromatograph (GC) and Agilent 7000C triple quadrupole mass spectrometer (MS/MS) with a J&W Select PAH column (30m x 0.25mm x 0.15 µm). Deuterated surrogates were used as internal standards for targets with similar physico-chemical properties. Instrument performance was verified at the beginning and end of each run and every 10–15 samples with a calibration check using Standard Reference Material (SRM) 1991 (National Institute of Standards and Technology). Solvent blanks were analyzed every 10–15 samples and at the beginning and end of each run. Duplicate sample aliquots were analyzed every 15 samples. In all cases, checked standards, matrix spikes, and duplicate samples met laboratory data quality objectives (within 30% of nominal or relative percent difference less than 30%).

### Microbial 16S rRNA library preparation and sequencing

Fecal samples were self-collected by human participants using the OMNIgene GUT OM200 Stool Sampling Kit (DNA Genotek Inc, Ottawa, Ontario, Canada) and shipped to the Microbiome Core Facility at Oregon State University for downstream processing. Stool samples were frozen at -80°C prior to processing all samples for DNA extraction and 16S rRNA library preparation in a single batch following our prior work (53). Briefly, after a 10-minute incubation, samples were subjected to bead beating. PCR was performed in triplicate using 1 microliter of purified DNA from the lysis solution to amplify the V4 region of the 16S rRNA gene, using the 806r and 515f primer set (54). An equal quantity of DNA was selected from each of the samples, and the pooled collection of DNA was cleaned then diluted and pooled prior to sequencing by the Center for Quantitative Life Sciences at Oregon State University, using a MiSeq instrument (Illumina, San Diego, California, USA) with 300-bp paired-end reads.

### PAH and Gut Microbiome Analyses

Using the targeted PAH exposure metabolite dataset, we investigated the relationship between PAH exposure to covariates including neurobehavioral group (HH/LL) and sex. Overdispersion was tested using a Poisson model of PAH abundance, evaluating PAHs and PAH metabolites which displayed a dispersion parameter greater than 1. Differences in PAH exposure abundance were modeled as a function of HH versus LL neurobehavioral group, with negative binomial generalized linear models used due to high overdispersion among nearly all metabolites. P-values from these models were adjusted using false-discovery rate, selecting only metabolites that were associated with neurobehavioral group below a pre-defined q-value of 0.1. We also implemented redundancy analysis to perform direct gradient analysis, creating a distance matrix based on PAH abundance using the Euclidean distance metric. We subsequently tested the relationship between neurobehavioral group and overall PAH exposure considering sex-dependent effects using PERMANOVA.

Gut microbiome sequences were filtered for read quality and clustered into amplicon sequence variants (ASVs) using DADA2 (55) with R 4.1.2 and taxonomically annotated using the SILVA reference annotation following our prior approach (56). Alpha- and beta-diversity analyses were performed using a relative abundance-normalized sequence count table. Generalized linear models were used to model species richness, phylogenetic richness, and Shannon diversity as a function of neurobehavioral group (HH vs. LL). Unweighted UniFrac ordination axes were generated using vegan (57). To clarify how neurobehavioral group relates to gut community composition, ANOVA was applied to the capscale-generated model of unweighted UniFrac distances.

Correlation between overall PAH exposure profile and the composition of the gut microbiome was assessed using a Mantel test with Spearman correlation (5,000 permutations), comparing pairwise Euclidean distances of standardized PAH concentrations against pairwise Bray-Curtis dissimilarities computed from rarefied 16S rRNA gene amplicon data. A linear model on the pairwise distance matrices was also fit as a parametric complement to the Mantel test.

To identify associations between individual microbial genera and PAH exposure, and between microbial genera and neurobehavioral group, we used MaAsLin2 (Microbiome Multivariable Associations with Linear Models, v1.22.0) (58). Genus-level abundances were total-sum-scaled (TSS) normalized and log-transformed prior to model fitting. For taxon–PAH associations, each genus was modeled as a function of scaled PAH concentration for each of 30 PAH compounds individually. For taxon–impulsivity associations, each genus was modeled as a function of cohort membership (HH vs. LL). P-values were adjusted using the Benjamini-Hochberg false discovery rate correction across all tests within each analysis, with significance defined at FDR < 0.1. Visualizations were created using ggplot2 (59).

## RESULTS

### PAH exposure is highly variable and overdispersed across the EMC2 cohort

Earlier work has demonstrated that PAH exposure is not uniform across human populations (60–63). Our findings are consistent with prior work testing the hypothesis that PAH exposure is highly variable and disproportionately experienced across cohort participants. PAHs vary in at least 10% of subjects, with more than 88% of the selected PAHs’ variance of exposure being much greater than the mean (GLM-Poisson, dispersion parameter > 1). This variation is readily observed visually, where several individuals display exposure to PAHs, such as naphthalene, at 2-3 orders of magnitude greater levels than other subjects in this data (Figure 1).

**FIG 1.**
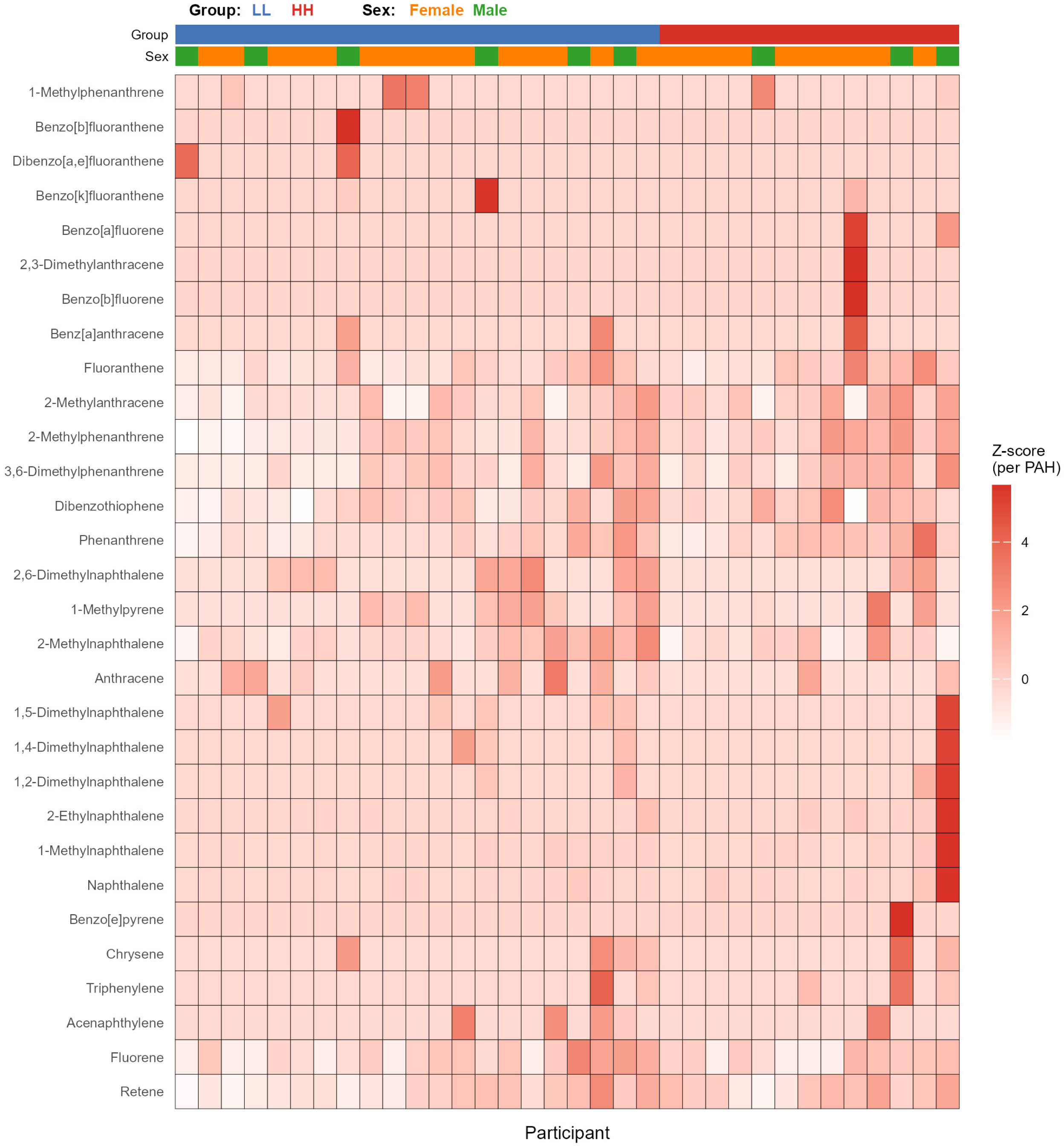
PAH exposure across the EMC2 cohort. Heatmap of 34 PAH compounds (rows) measured from silicone wristbands worn by 34 participants (columns). Values are Z-score normalized within each PAH compound, with darker shading indicating higher relative exposure. Participants are grouped by neurobehavioral classification (LL = low impulsivity/fast attention, blue; HH = high impulsivity/poor attention, red) and sex (female, orange; male, green), as indicated by the annotation bars.

### Impulsivity and attention are linked to PAH exposure in a sex-dependent fashion

Previous work has shown that exposure to PAHs and other xenobiotics is correlated with behaviors which are related to the cognitive trait of impulsivity and attention problems (13, 16, 17). We explored the hypothesis that PAH exposure similarly links to impulsivity and attention in the EMC2 cohort. Our results uncover a significant association between neurobehavioral group membership and exposure to specific PAHs. Individuals from the HH group exhibited markedly elevated levels of four PAHs: naphthalene (q = 0.009), 2-ethylnaphthalene (q = 0.035), 1-methylnaphthalene (q = 0.035), and 2-methylphenanthrene (q = 0.066; GLM; Table S4; Figure 2a). No PAHs were enriched in the LL group.

**FIG 2.**
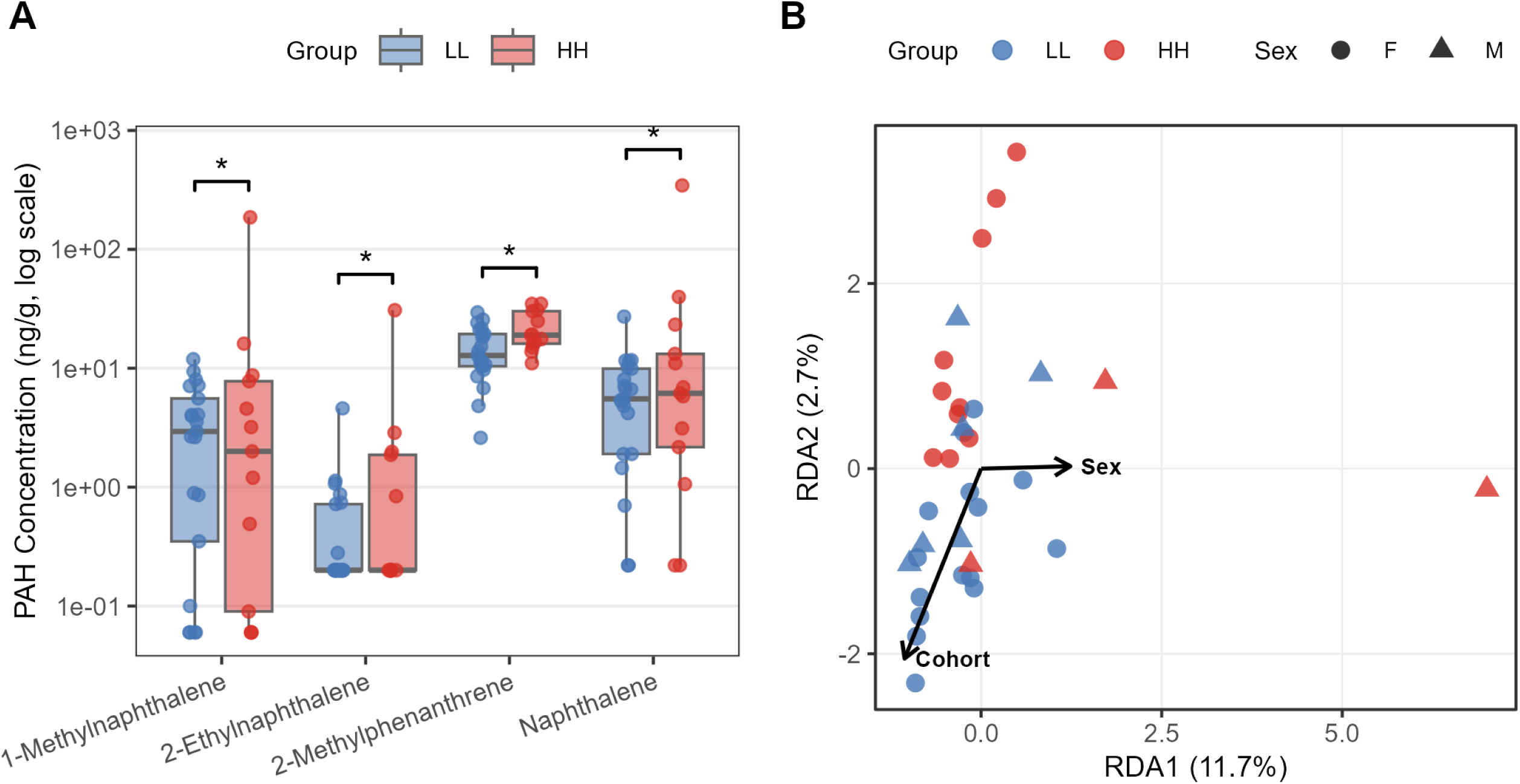
PAH exposure differs by neurobehavioral group and sex. (A) PAH concentrations (ng/g, log scale) for the four compounds significantly associated with neurobehavioral group (negative binomial generalized linear model, FDR < 0.1). HH participants (high impulsivity/poor attention, red) exhibited elevated levels of naphthalene, 1-methylnaphthalene, 2-ethylnaphthalene, and 2-methylphenanthrene relative to LL participants (low impulsivity/fast attention, blue). Asterisks denote FDR < 0.1. (B) Redundancy analysis (RDA) of overall PAH exposure profiles constrained by neurobehavioral group and sex. Points represent individual participants colored by group (LL, blue; HH, red) and shaped by sex (circle, female; triangle, male). Vectors indicate the direction and relative magnitude of cohort (HH/LL) and sex effects. A significant interaction between sex and neurobehavioral group predicted PAH exposure composition (PERMANOVA, p = 0.026).

Variation in the metabolism of PAHs has been associated with factors ranging from host-genomic capacity to the metabolic activity of the gut microbiome, and it has been shown that PAH detoxification may result in increased production of estrogenic metabolites (28) and sex-dependent factors that influence PAH detoxification (18, 19, 64, 65). Sex-dependent differences are reported for impulsivity (66–68); therefore, a redundancy analysis explored the potential relationship between sex and the overall impulsivity-PAH connection. A significant interaction between sex and neurobehavioral group predicted PAH exposure (PERMANOVA, p = 0.026, Figure 2b). Males exhibited higher PAH levels than females, with this disparity most pronounced in HH males. This finding suggests a potential sex-dependent modulation of the neurobehavioral-PAH association and may reflect underlying biological or behavioral mechanisms that differentially regulate a participant’s exposure to PAHs.

### Gut microbiome diversity is linked to neurobehavioral group and PAH exposure

Variation in the microbiome has been linked to exposure to specific PAHs in prior work (28, 34, 35); therefore, we sought to determine if the composition of the gut microbiome links to the overall PAH exposure profile an individual experiences. Using a Mantel correlation test to examine this relationship, we find positive correlation between PAH exposure distance (Euclidean distance) and Bray-Curtis microbiome dissimilarity (Mantel test, r = 0.21, p = 0.034). Moreover, a linear model that uses pairwise Euclidean distances between subjects based on PAH exposure to predict pairwise Bray-Curtis dissimilarity suggests a positive correlation between these metrics of PAH exposure and microbiome diversity (linear model, slope = 8.49, p < 0.0001, R² = 0.057; Supplementary Figure S1). Collectively, these results highlight that variability in PAH exposure tracks with variability in microbiome composition, underscoring the potential for exposure profiles to contribute to microbiome individuality.

We next sought to determine if variation in the gut microbiome also links to neurobehavioral group membership. We evaluated this relationship using linear models and found that participants in the HH group possessed microbiomes that were more taxonomically and phylogenetically diverse. Specifically, HH group membership was statistically associated with larger levels of microbiome taxonomic richness and Shannon entropy, as well as phylogenetic diversity (Generalized Linear Model; observed richness p = 0.033, Shannon p = 0.048, Faith’s PD p = 0.016; Figure 3a). We also evaluated how the overall composition of the gut microbiome associated with neurobehavioral group, finding a statistically significant relationship between microbiome unweighted-UniFrac distance and HH/LL membership (capscale ANOVA, p=0.038, Figure 3b). Collectively, these results indicate that the gut microbiome correlates with neurobehavioral phenotype at the community level, and points to a relationship between PAH exposure, neurobehavioral function, and the gut microbiome that warrants further investigation.

**FIG 3.**
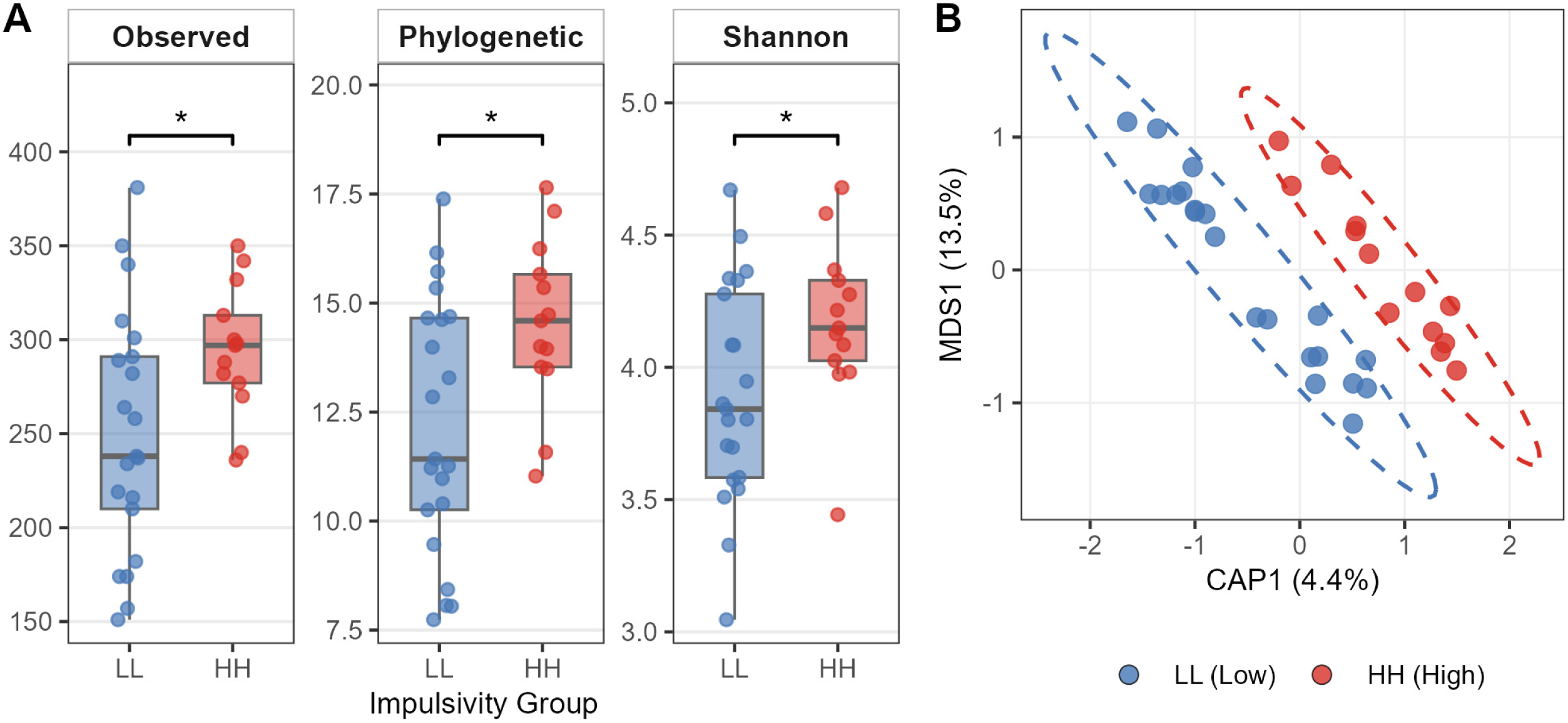
Gut microbiome diversity differs by neurobehavioral group. (A) Alpha-diversity metrics — observed taxonomic richness, Faith’s phylogenetic diversity, and Shannon entropy — compared between LL (blue) and HH (red) participants. HH group membership was associated with significantly higher values for all three metrics (generalized linear model; observed richness p = 0.033, phylogenetic diversity p = 0.016, Shannon p = 0.048). Asterisks denote p < 0.05. (B) Constrained ordination (capscale) of gut microbiome composition using unweighted UniFrac distances. Points represent individual participants colored by neurobehavioral group, with dashed ellipses indicating group dispersion. Neurobehavioral group was significantly associated with community composition (capscale ANOVA, p = 0.038).

### Specific microbial genera are associated with PAH exposure

We next sought to determine if specific taxa may underlie the community-level relationship between PAH exposure and microbiome variation. Sequences were agglomerated to the genus level, and MaAsLin2 was used to test associations between genus-level abundance and individual PAH compound concentrations. Of 5,160 genus-PAH pairs tested, 33 exhibited significant associations after correction for multiple testing (FDR < 0.1), involving 21 unique genera and 14 PAH compounds (Table S5). The strongest association was observed between *Prevotella* and benzo[b]fluoranthene, followed by *UC5-1-2E3* and benzo[b]fluoranthene. *Haemophilus* and *Lactobacillus* both exhibited significant positive associations with 1-methylphenanthrene, while *Oxalobacter* and *Papillibacter* were associated with benzo[e]pyrene. *Erysipelatoclostridium* showed a significant positive association with anthracene, and a *Ruminococcaceae* member was positively associated with 2,6-dimethylnaphthalene. Three genera showed negative associations with PAH exposure: *Bacteroides* with benzo[b]fluoranthene, *Dorea* with 1,4-dimethylnaphthalene, and *Lachnospiraceae* UCG-004 with acenaphthylene, each exhibiting decreased abundance with higher PAH exposure. Overall, 30 of 33 significant associations were positive, indicating that higher PAH exposure generally tracked with increased genus-level relative abundance (Figure 4).

**FIG 4.**
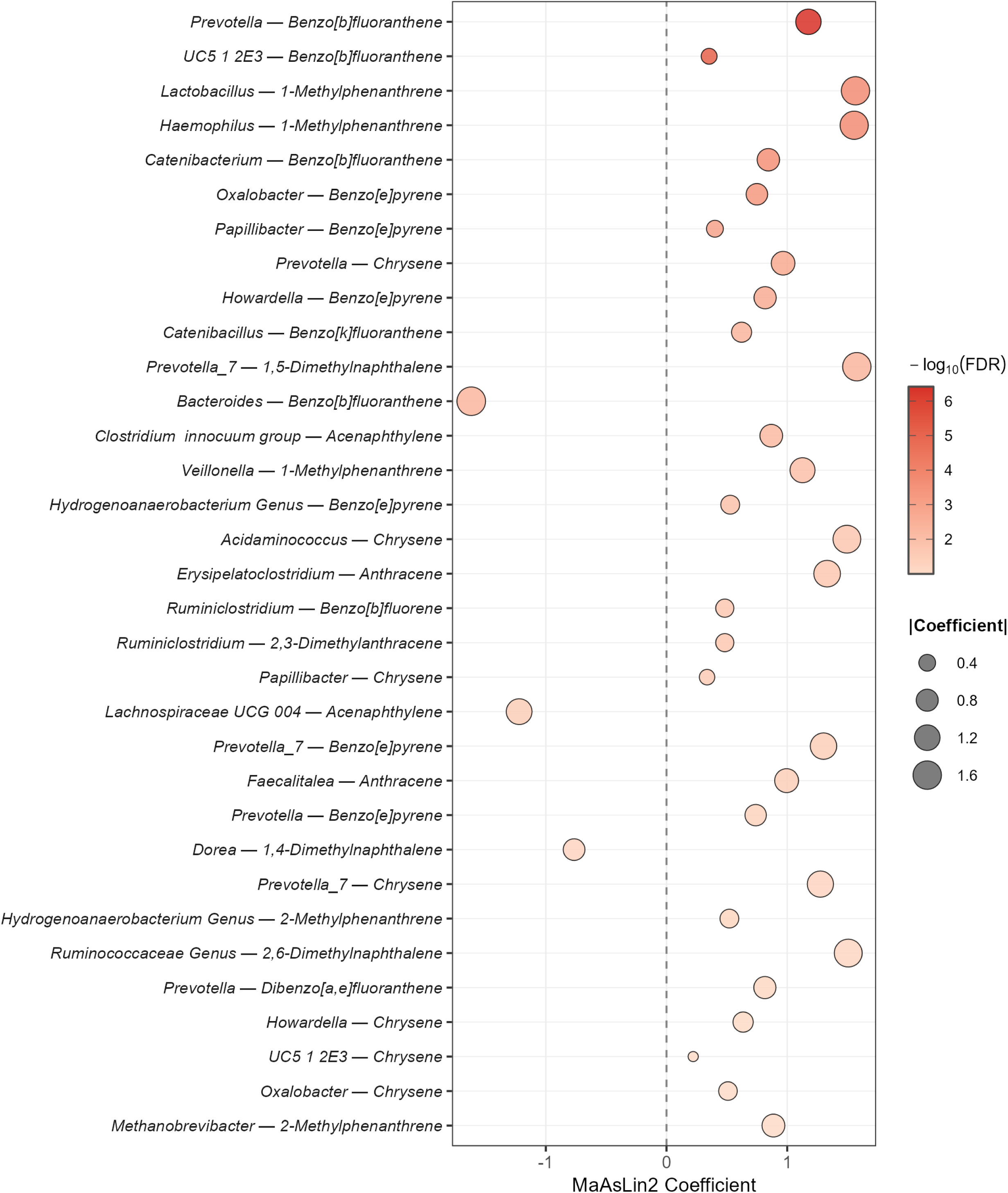
Genus-level associations with PAH exposure identified by MaAsLin2. Each point represents a significant association (FDR < 0.1) between a microbial genus and an individual PAH compound. The x-axis shows the MaAsLin2 coefficient (positive values indicate increased genus abundance with higher PAH exposure; negative values indicate decreased abundance). Point size reflects the absolute value of the coefficient, and color intensity reflects statistical significance (−log₁₀ FDR). Of 33 significant associations involving 21 genera and 14 PAH compounds, 30 were positive, indicating that higher PAH exposure generally tracked with increased genus-level relative abundance. Three genera — *Bacteroides*, *Dorea*, and *Lachnospiraceae* UCG-004 — showed negative associations with PAH exposure.

No individual genus reached significance for differential abundance between HH (high impulsivity/poor attention, n=13) and LL (low impulsivity/fast attention, n=21) participants after correction for multiple testing (minimum FDR = 0.48). Among the 172 genera tested, 14 showed nominal associations with neurobehavioral group (p < 0.05; Table S6, Figure S2). Thirteen of these were more abundant in HH participants, including *Collinsella* (p = 0.007), *Angelakisella* (p = 0.008), *Hydrogenoanaerobacterium* (p = 0.018), and *Methanobrevibacter* (p = 0.031); *Tuzzerella* was the sole genus with higher LL abundance (p = 0.028). This directional consistency, with nearly all nominally associated taxa enriched in the high-impulsivity/poor-attention group, parallels the alpha-diversity finding of greater taxonomic richness in HH participants. The absence of individually significant taxa after multiple testing correction is consistent with the limited statistical power of this pilot study, which affords approximately 1% power to detect medium effect sizes (d = 0.5) across 172 tests at n = 34. Nevertheless, the 14 nominally associated genera provide a candidate set for cross-referencing with the PAH-responsive taxa identified above.

## DISCUSSION

Prior studies independently link PAH exposure to the gut microbiome (28, 34, 35), PAH exposure to impulsivity and attention (13–17), and the microbiome to neurobehavioral variation (69–72), but these three factors have not previously been examined together in a single human cohort. Here, we demonstrate that PAH exposure, gut microbiome composition, and neurobehavioral function covary within a geographically distributed Oregon population. We observed a consistent association between cumulative PAH exposure and neurobehavioral group, modulated by sex, and both variables were independently linked to shifts in gut microbiome composition at the community level. Overall PAH mixture profiles correlated with microbiome dissimilarity, and neurobehavioral group associated with both alpha- and beta-diversity of the gut microbiome. At the taxon level, 21 microbial genera were significantly associated with specific PAH compounds, and a subset of these overlapped with genera nominally associated with neurobehavioral group membership. Together, these results support a three-way relationship among environmental chemical exposure, gut microbiome composition, and neurobehavioral function, and identify candidate taxa at their intersection for targeted investigation.

The relationship between PAH exposure and neurobehavioral phenotype was moderated by sex, with the disparity between HH and LL groups most pronounced among males. This observation is consistent with prior work showing notable differences in the metabolism of PAHs between males and females (18, 19, 64, 65), and with evidence that cell lines from male and female subjects show extensive differences in the abundance of gene transcripts directly related to PAH exposure (73, 74). Several mechanisms could underlie this sex-dependent relationship. Byproducts of PAH metabolism are known to disrupt endocrine signaling, particularly androgen hormone signaling (75, 76), and PAH compounds and their metabolic byproducts associate with hormone levels in human studies (19, 77, 78). Changes in hormone signaling can have profound impacts on cognitive function, including impulsivity and attention (79, 80). However, it is important to note that our data establish a statistical interaction between sex and neurobehavioral group in predicting PAH exposure; the mechanistic chain from PAH metabolism through endocrine disruption to behavioral change involves several inferential steps that remain to be tested directly.

Beyond the PAH-behavior link, PAH exposure was also associated with gut microbiome composition at the community level. The significant Mantel correlation between pairwise PAH Euclidean distance and Bray-Curtis microbiome dissimilarity (r = 0.21, p = 0.034) indicates that participants with more similar PAH exposure profiles tend to harbor more similar gut communities. While the effect size is modest (R² = 0.057 in the complementary linear model), it is notable in a cohort of this size and consistent with the hypothesis that PAH mixtures – not just individual compounds – exert selective pressure on gut community composition. This finding aligns with animal model studies demonstrating that PAH exposure reshapes overall gut community structure (32–35) and extends these observations to a free-living human population experiencing environmentally relevant, real-world exposure levels.

The relationship between neurobehavioral function and the gut microbiome is an area of active investigation, but behaviors associated with impulsivity, including smoking (81), excessive alcohol consumption (82), and increased consumption of processed carbohydrates (83), have all been linked to changes in the diversity of the gut microbiome. The present work provides evidence that neurobehavioral group and PAH exposure are linked to the diversity and composition of the gut microbiome at the community level. HH group membership was associated with increased phylogenetic and taxonomic richness, and the overall composition of the gut microbiome differed between HH and LL participants as measured by unweighted UniFrac distance. The nature of this diversity difference is informative: the fact that HH participants showed greater taxonomic richness alongside significant unweighted UniFrac separation, a metric sensitive to the presence or absence of rare lineages, suggests that neurobehaviorally associated microbiome variation involves the gain of less common taxa rather than shifts in dominant community members. The direction of this relationship is not certain, but it is possible that individuals with higher impulsivity are exposed to greater microbial diversity through their diets or environments. Impulse-related behaviors such as smoking and excessive alcohol consumption can disrupt microbial community dynamics by altering immune system homeostasis (84, 85) and increasing toxic chemical exposure (86), potentially dysregulating immune recognition of commensal taxa and allowing proliferation of transient populations. These behavioral mechanisms offer a plausible route by which the impulsivity component of HH group membership could contribute to the observed microbiome differences, though the independent contribution of attention deficits to gut microbiome variation remains unexplored.

Conversely, the gut microbiome may impact variation in neurobehavioral function. Gut microbiome composition is an established driver of serotonin physiology (87–90), and serotonin is a suggested modulator of impulsivity, including sex-dependent differences (91). Gut microbes also produce a range of neuroactive metabolites, including short-chain fatty acids and neurotransmitter precursors, that influence neural function through immune, endocrine, and vagal pathways (39–41). These microbiome-brain pathways suggest at least two routes through which PAH exposure could alter neurobehavioral function via the gut microbiome. First, PAH-induced disruption of microbial community structure could destabilize serotonin-related host-microbe interactions, indirectly altering serotonergic signaling to the brain. Second, microbial biotransformation of PAHs could generate estrogenic metabolites that differentially affect males and females, producing sex-divergent behavioral outcomes. This route is supported by evidence that human intestinal microbiota convert PAHs into potent estrogen receptor activators (28) and is consistent with the sex-dependent PAH-neurobehavioral relationship observed in this cohort. Distinguishing between these pathways will require metabolomic profiling of PAH-derived microbial metabolites in stool, coupled with characterization of circulating estrogen receptor agonist concentrations and serotonergic markers, in cohorts large enough to stratify by sex.

Among the 21 PAH-responsive genera, several have established biological roles that provide context for their sensitivity to PAH exposure. *Prevotella*, associated with high-fiber, plant-rich diets and carbohydrate fermentation (92, 93), showed significant positive associations with benzo[b]fluoranthene, benzo[e]pyrene, chrysene, and dibenzo[a,e]fluoranthene, while *Prevotella_7*, a phylogenetically distinct clade in the SILVA taxonomy, was independently associated with benzo[e]pyrene, chrysene, and 1,5-dimethylnaphthalene. *Lactobacillus*, which showed a significant association with 1-methylphenanthrene, has well-characterized effects on gut-brain axis signaling, including modulation of stress responses and neurotransmitter precursor production (94). Three genera showed negative PAH associations, including *Bacteroides* (benzo[b]fluoranthene, FDR = 0.040), *Dorea* (1,4-dimethylnaphthalene), and *Lachnospiraceae* UCG-004 (acenaphthylene). *Bacteroides* is a dominant commensal whose depletion with PAH exposure could have functional consequences given its central roles in polysaccharide metabolism and immune regulation (95). While these taxon-level associations establish which genera respond to individual PAH compounds, a key question is whether any of these PAH-responsive taxa also track with neurobehavioral phenotype.

Individual taxa did not reach significance for differential abundance between neurobehavioral groups after multiple testing correction, consistent with the limited statistical power of this pilot cohort. However, a central question motivating this study is whether specific gut taxa sit at the intersection of PAH exposure and neurobehavioral phenotype. Cross-referencing the 21 genera with significant PAH associations (FDR < 0.1) against the 14 genera with nominal neurobehavioral associations (p < 0.05) identified two genera with directionally coherent associations across both domains. *Hydrogenoanaerobacterium* showed significant positive associations with benzo[e]pyrene (FDR = 0.013) and 2-methylphenanthrene (FDR = 0.065) and was nominally enriched in HH participants (p = 0.018). *Methanobrevibacter* showed a significant positive association with 2-methylphenanthrene (FDR = 0.099) and was nominally enriched in HH participants (p = 0.031). For both genera, higher PAH exposure and HH group membership associate with greater abundance. Independent literature supports both as biologically plausible candidates. *Hydrogenoanaerobacterium* abundance has been negatively correlated with memory scores in mild cognitive impairment patients (96) and linked to Parkinson’s disease (97). *Methanobrevibacter* is enriched in Parkinson’s disease patients with impulse control disorders, with concurrent enrichment of xenobiotic degradation pathways (97) and is associated with cognitive impairment severity in aged care populations (98) and neuroinflammatory signaling in multiple sclerosis (99, 100). Targeted studies of these candidates in larger cohorts, especially those that incorporate longitudinal designs, metagenomic and metabolomic profiling, and temporal ordering of exposure, microbiome change, and behavioral outcomes, represent a direct path toward testing whether the microbiome mediates the relationship between environmental PAH exposure and neurobehavioral phenotypes.

As a pilot study, this work establishes the empirical basis for several targeted extensions. Larger cohorts with complete paired data will be needed to move beyond the community-level associations detected here (n = 34) and to robustly identify genus-level associations across the full taxonomic breadth after multiple testing correction. Longitudinal sampling designs can address what the present cross-sectional design cannot: the temporal ordering of PAH exposure, microbiome change, and behavioral outcomes necessary to evaluate causal directionality. Our 16S rRNA amplicon approach characterizes which taxa respond to PAH exposure, but metagenomic and metabolomic profiling will be essential to determine whether these taxa are actively metabolizing PAHs or producing neuroactive metabolites. Similarly, integrating longer-term exposure assessment beyond the 30-day wristband window used here with repeated microbiome sampling would clarify whether the associations we observe reflect acute responses or stable, exposure-shaped community states. Together, these extensions would provide the multi-omic depth and temporal resolution needed to test whether the gut microbiome mediates the link between environmental PAH exposure and neurobehavioral outcomes.

Collectively, our results provide evidence that PAH exposure, gut microbiome composition, and neurobehavioral function covary within a single human cohort, with associations observable at the community level and, for PAH exposure, at the level of specific microbial genera. The sex-dependent modulation of the exposure–behavior relationship is consistent with prior work documenting sex differences in PAH metabolism and suggests that endocrine-mediated pathways may shape how environmental exposures translate into neurobehavioral variation. By demonstrating that all three domains converge within the same population, this work moves beyond pairwise associations to establish an empirical foundation for testing whether the gut microbiome mediates the link between environmental chemical exposure and neurobehavioral outcomes.

## ACKNOWLEDGMENTS

The EMC2 study was funded by the Pacific Northwest Center for Translational Environmental Health Research, a P30 Center funded by the National Institutes of Environmental Health Sciences (P30ES030287). HOP research infrastructure was funded by the Cancer Early Detection Advanced Research Center at Oregon Health & Science University’s Knight Cancer Institute and the National Cancer Institute (P30CA069533 and 1U01CA232819 to JS). OHSU’s Oregon Clinical and Translational Research Institute (OCTRI; 1UL1TR002369) developed technology infrastructure securely linking the HOP app with *Let’s Get Healthy!* modules used for data collection and e-feedback, including neurobehavioral measures in this study. *Let’s Get Healthy!* was developed through funding from the National Institutes of Health (NIH), including Science Education Partnership Awards (R25OD01496, R25GM129840 to LKM) and NIH grants (R25RR020443-05S1, UL1RR024140-04S3, RR026008, 3P30CA69553-13S9, and UL1TR002369).

## DATA AVAILABILITY

The data underlying the findings of this study are available in the NCBI Sequence Read Archive (SRA) under BioProject ID PRJNA1344322.

## Supplemental Materials

**TABLE S1.** Demographic and sample characteristics of EMC2 participants with complete paired data (n = 34). Participants were classified into HH (high impulsivity/poor attention; n = 13) and LL (low impulsivity/fast attention; n = 21) neurobehavioral groups. Kit transit time refers to the number of days between kit delivery to the participant and return via mail. P-values are from two-sample t-tests (continuous variables) or Fisher’s exact test (sex). Groups did not differ significantly in age, sex distribution, kit transit time, or 16S rRNA sequencing depth.

**TABLE S2.** Cross-tabulation of the EMC2 recruitment pool by impulsivity quartile (BIS-15 total score) and attention quartile (PVT-B median reaction time) among 2,628 participants who completed both assessments. Highlighted cells denote the LL recruitment group (Q1 impulsivity × Q1 attention; low impulsivity, fast reaction times) and the HH recruitment group (Q4 impulsivity × Q4 attention; high impulsivity, slow reaction times). Group sizes did not differ significantly (Chi-square p = 0.33). Note: missing closing parentheses in Q4 row total and Q3 column total should be corrected.

**TABLE S3.** Attention metrics differ between lowest (Q1) and highest (Q4) impulsivity quartiles. Independent-samples t-tests comparing median reaction time, lapses (errors of omission), and false starts (errors of commission) between Q1 (n = 283) and Q4 (n = 139) participants. All metrics differed significantly (p < 0.005). Levene’s test indicated unequal variances for all metrics (p < 0.002); degrees of freedom and test statistics reflect Welch’s correction.

**TABLE S4.** Negative binomial generalized linear model results for PAH compound concentrations as a function of neurobehavioral group (HH vs. LL). Each of 30 PAH compounds detected in silicone wristbands was modeled independently. Columns show group means (ng/g), model estimate, standard error, z-statistic, uncorrected p-value, Benjamini-Hochberg FDR-adjusted q-value, and direction of enrichment. Four PAHs were significantly elevated in HH participants after multiple testing correction (FDR < 0.1): naphthalene, 1-methylnaphthalene, 2-ethylnaphthalene, and 2-methylphenanthrene. No PAHs were significantly enriched in the LL group. Results are sorted by FDR.

**TABLE S5.** MaAsLin2 results for all significant genus–PAH associations (FDR < 0.1). All 33 significant associations involving 21 unique genera and 14 PAH compounds are listed with effect estimates, standard errors, p-values, and FDR-adjusted q-values. Genus-level abundances were TSS-normalized and log-transformed prior to model fitting.

**TABLE S6.** MaAsLin2 results for genus–neurobehavioral group associations. No genera reached significance after FDR correction (minimum FDR = 0.48). The 14 genera with nominal associations (uncorrected p < 0.05) are shown with effect estimates, standard errors, p-values, and FDR-adjusted q-values. Thirteen of 14 nominally associated genera were more abundant in HH participants.

**FIG S1.** Community-level correlation between PAH exposure and gut microbiome composition. Scatterplot of pairwise Bray-Curtis dissimilarity (x-axis; computed from rarefied 16S rRNA genus-level abundances) against pairwise Euclidean distance of standardized PAH concentrations (y-axis) for all participant pairs (n = 561 pairwise comparisons). Blue line indicates linear model fit (slope = 8.49, p < 0.0001, R² = 0.057); shaded band indicates 95% confidence interval. The Mantel test confirmed a significant positive correlation between PAH exposure distance and microbiome dissimilarity (Spearman r = 0.21, p = 0.034, 5,000 permutations).

**FIG S2.** Rarefied abundances of the 14 genera with nominal associations (uncorrected p < 0.05) with neurobehavioral group (HH vs. LL), identified by MaAsLin2. Each panel shows one genus, with individual data points overlaid on boxplots for LL (blue) and HH (red) participants. Thirteen of 14 genera were more abundant in HH participants; *Tuzzerella* was the sole genus with higher LL abundance. No genera reached significance after FDR correction (minimum FDR = 0.48).

## REFERENCES

1. Masclet P, Mouvier G, Nikolaou K. 1986. Relative decay index and sources of polycyclic aromatic hydrocarbons. Atmospheric Environment (1967) 20:439–446.

2. Mojiri A, Zhou JL, Ohashi A, Ozaki N, Kindaichi T. 2019. Comprehensive review of polycyclic aromatic hydrocarbons in water sources, their effects and treatments. Science of The Total Environment 696:133971.

3. Wilcke W. 2007. Global patterns of polycyclic aromatic hydrocarbons (PAHs) in soil. Geoderma 141:157–166.

4. Meador JP, Stein JE, Reichert WL, Varanasi U. 1995. Bioaccumulation of polycyclic aromatic hydrocarbons by marine organisms. Reviews of Environmental Contamination and Toxicology 143:79–165.

5. Ramesh A, Archibong AE, Hood DB, Guo Z, Loganathan BG. 2004. Bioavailability and risk assessment of orally ingested polycyclic aromatic hydrocarbons. International Journal of Toxicology 23:301–333.

6. Ramesh A, Archibong A, Hood D, Guo Z, Loganathan B. 2012. Global environmental distribution and human health effects of polycyclic aromatic hydrocarbons, p. 97–128. In Global Contamination Trends of Persistent Organic Chemicals. CRC Press.

7. 2003. PAHs: An ecotoxicological perspective. John Wiley & Sons.

8. Yan T, Chengzhi C, Haiyan Y, Baijie T. 2010. Distribution of benzo[a]pyrene in discrete regions of rat brain tissue using light microscopic autoradiography and gamma counting. Toxicological & Environmental Chemistry 92:1309–1317.

9. Khan A, Jahego AR. 2021. Polycyclic aromatic hydrocarbons and neurological disorders: From exposure to preventive interventions, p. 335–353. In Akash, MSH, Rehman, K (eds.), Environmental Contaminants and Neurological Disorders. Springer International Publishing, Cham.

10. Aschner M, Costa LG. 2015. Environmental factors in neurodevelopmental and neurodegenerative disorders. Elsevier Inc.

11. Patri M, Padmini A, Babu P. 2009. Polycyclic aromatic hydrocarbons in air and their neurotoxic potency in association with oxidative stress: A brief perspective. Annals of Neurosciences.

12. Perera FP, Tang D, Wang S, Vishnevetsky J, Zhang B, Diaz D, Camann D, Rauh V. 2012. Prenatal polycyclic aromatic hydrocarbon (PAH) exposure and child behavior at age 6-7 years. Environ Health Perspect 120:921–926.

13. Perera FP, Chang HW, Tang D, Roen EL, Herbstman J, Margolis A, Huang TJ, Miller RL, Wang S, Rauh V. 2014. Early-life exposure to polycyclic aromatic hydrocarbons and ADHD behavior problems. PLoS One 9:e111670.

14. Peterson BS, Rauh VA, Bansal R, Hao X, Toth Z, Nati G, Walsh K, Miller RL, Arias F, Semanek D, Perera F. 2015. Effects of prenatal exposure to air pollutants (polycyclic aromatic hydrocarbons) on the development of brain white matter, cognition, and behavior in later childhood. JAMA Psychiatry 72:531–540.

15. Mortamais M, Pujol J, Martínez-Vilavella G, Fenoll R, Reynes C, Sabatier R, Rivas I, Forns J, Vilor-Tejedor N, Alemany S, Cirach M, Alvarez-Pedrerol M, Nieuwenhuijsen M, Sunyer J. 2017. Effect of exposure to polycyclic aromatic hydrocarbons on basal ganglia and attention-deficit hyperactivity disorder symptoms in primary school children. Environment International 105:12–19.

16. Perez-Fernandez C, Flores P, Sánchez-Santed F. 2019. A systematic review on the influences of neurotoxicological xenobiotic compounds on inhibitory control. Frontiers in Behavioral Neuroscience 13:139.

17. Bourque J, Mendrek A, Dinh-Williams L, Potvin S. 2013. Neural circuitry of impulsivity in a cigarette craving paradigm. Frontiers in Psychiatry 4.

18. Zhao C, Zhu L, Li J, Zhou H, Chen Y, Wang T, Luo J, Ma S, Pei S, Zhu Y, Zhang Y. 2022. Parent and halogenated polycyclic aromatic hydrocarbons in the serum of coal-fired power plant workers: Levels, sex differences, accumulation trends, and risks. Environmental Science & Technology 56:12431–12439.

19. Yuan Q, Luo K, Zhang P, Liu B, Chen Y, Ding Z. 2022. Urinary polycyclic aromatic hydrocarbon metabolites are positively related to serum testosterone levels of males and serum estradiol levels of females among U.S. adults. Frontiers in Endocrinology 13.

20. Schroeder H. 2011. Developmental brain and behavior toxicity of air pollutants: a focus on the effects of polycyclic aromatic hydrocarbons (PAHs). Critical Reviews in Environmental Science and Technology 41:2026–2047.

21. Olasehinde TA, Olaniran AO. 2022. Neurotoxicity of polycyclic aromatic hydrocarbons: a systematic mapping and review of neuropathological mechanisms. Toxics 10:417.

22. Claus SP, Guillou H, Ellero-Simatos S. 2016. The gut microbiota: a major player in the toxicity of environmental pollutants? NPJ Biofilms and Microbiomes 2:16003.

23. Sutherland VL, McQueen CA, Mendrick D, Gulezian D, Cerniglia C, Foley S, Forry S, Khare S, Liang X, Manautou JE, Tweedie D, Young H, Alekseyenko AV, Burns F, Dietert R, Wilson A, Chen C. 2020. The Gut Microbiome and Xenobiotics: Identifying Knowledge Gaps. Toxicological Sciences 176:1–10.

24. Sowada J, Schmalenberger A, Ebner I, Luch A, Tralau T. 2014. Degradation of benzo[a]pyrene by bacterial isolates from human skin. FEMS Microbiology Ecology 88:129–139.

25. Cerniglia CE, Sutherland JB. 2010. Degradation of polycyclic aromatic hydrocarbons by fungi, p. 2079–2110. In Timmis, KN (ed.), Handbook of Hydrocarbon and Lipid Microbiology. Springer, Berlin, Heidelberg.

26. Lemoine L, Gonzalez-Mariscal I, Alayrac P, Dairou J, Coumoul X, Barouki R, Kerdine-Römer S, Langouët S. 2021. Commensal-related changes in the epidermal barrier function lead to alterations in the benzo[a]pyrene metabolite profile and its distribution in 3D skin. mBio 12:e01223–21.

27. Sowada J, Lemoine L, Schön K, Hutzler C, Luch A, Tralau T. 2017. Toxification of polycyclic aromatic hydrocarbons by commensal bacteria from human skin. Archives of Toxicology 91:2331–2341.

28. Van de Wiele T, Vanhaecke L, Boeckaert C, Peru K, Headley J, Verstraete W, Siciliano S. 2005. Human colon microbiota transform polycyclic aromatic hydrocarbons to estrogenic metabolites. Environmental Health Perspectives 113:6–10.

29. Aris AZ, Shamsuddin AS, Praveena SM. 2014. Occurrence of 17α-ethynylestradiol (EE2) in the environment and effect on exposed biota: a review. Environment International 69:104–119.

30. Zhang H, Chen J, Li Z, Liu L. 2021. Testing for mediation effect with application to human microbiome data. Statistics in Biosciences 13:313–328.

31. Chiu K, Warner G, Nowak RA, Flaws JA, Mei W. 2020. The impact of environmental chemicals on the gut microbiome. Toxicological Sciences 176:253–284.

32. Du B, Zhang D, Liu Y, Zhou G, Pan C, Guo G, Yang K, Zhao G, Zhang J. 2023. Evaluation of the impact of BaP exposure on the gut microbiota and allergic responses in an OVA-sensitized mouse model. Environmental Health Perspectives 131:067004.

33. Ribière C, Peyret P, Parisot N, Darcha C, Déchelotte PJ, Barnich N, Peyretaillade E, Boucher D. 2016. Oral exposure to environmental pollutant benzo[a]pyrene impacts the intestinal epithelium and induces gut microbial shifts in murine model. 1. Sci Rep 6:31027.

34. Xie S, Zhou A, Wei T, Li S, Yang B, Xu G, Zou J. 2020. Benzo[a]pyrene induces microbiome dysbiosis and inflammation in the intestinal tracts of western mosquitofish (Gambusia affinis) and zebrafish (Danio rerio). Fish & Shellfish Immunology 105:24–34.

35. Stagaman K, Alexiev A, Sieler MJ, Hammer A, Kasschau KD, Truong L, Tanguay RL, Sharpton TJ. 2024. The zebrafish gut microbiome influences benzo[a]pyrene developmental neurobehavioral toxicity. Sci Rep 14:14618.

36. Messaoudi M, Violle N, Bisson JF, Desor D, Javelot H, Rougeot C. 2011. Beneficial psychological effects of a probiotic formulation (Lactobacillus helveticus R0052 and Bifidobacterium longum R0175) in healthy human volunteers. Gut Microbes 2:256–261.

37. Akkasheh G, Kashani-Poor Z, Tajabadi-Ebrahimi M, Jafari P, Akbari H, Taghizadeh M, Memarzadeh MR, Asemi Z, Esmaillzadeh A. 2016. Clinical and metabolic response to probiotic administration in patients with major depressive disorder: A randomized, double-blind, placebo-controlled trial. Nutrition 32:315–320.

38. Steenbergen L, Sellaro R, van Hemert S, Bosch JA, Colzato LS. 2015. A randomized controlled trial to test the effect of multispecies probiotics on cognitive reactivity to sad mood. Brain, Behavior, and Immunity 48:258–264.

39. Ahmed H, Leyrolle Q, Koistinen V, Kärkkäinen O, Layé S, Delzenne N, Hanhineva K. 2022. Microbiota-derived metabolites as drivers of gut–brain communication. Gut Microbes 14:2102878.

40. Appleton J. 2018. The Gut-Brain Axis: Influence of Microbiota on Mood and Mental Health. Integr Med (Encinitas) 17:28–32.

41. Chakrabarti A, Geurts L, Hoyles L, Iozzo P, Kraneveld AD, La Fata G, Miani M, Patterson E, Pot B, Shortt C, Vauzour D. 2022. The microbiota–gut–brain axis: pathways to better brain health. Perspectives on what we know, what we need to investigate and how to put knowledge into practice. Cellular and Molecular Life Sciences 79:80.

42. O’Brien TD, Potter AB, Driscoll CC, Goh G, Letaw JH, McCabe S, Thanner J, Kulkarni A, Wong R, Medica S, Week T. 2023. Population screening shows risk of inherited cancer and familial hypercholesterolemia in Oregon. The American Journal of Human Genetics 110:1249–1265.

43. Zhang Z, Shafer A, Johnson-Camacho K, Adey A, Anur P, Brown KA, Shannon J. 2024. Novel recruitment approaches and operational results for a statewide population cohort for cancer research: The Healthy Oregon Project. Journal of Clinical and Translational Science 8:e32.

44. Spinella M. 2007. Normative data and a short form of the Barratt Impulsiveness Scale. International Journal of Neuroscience 117:359–368.

45. Meule A. 2023. Cut-off scores for the Barratt Impulsiveness Scale–short form (BIS–15): sense and nonsense. International Journal of Neuroscience 134:1149–1152.

46. Juneja R, Chaiwong W, Siripool P, Mahapol K, Wiriya T, Shannon JS, Petchkrua W, Kunanusont C, Marriott LK. 2019. Thai adaptation and reliability of three versions of the Barratt Impulsiveness Scale (BIS 11, BIS-15, and BIS-Brief). Psychiatry Research 272:744–755.

47. Marriott LK, Coppola L, Mitchell SH, Bouwma-Gearhart J, Zhen Z, Shifrer D, Feryn AB, Shannon J. 2019. Opposing effects of impulsivity and mindset on science self-efficacy and STEM interest in adolescents. PLOS One 14:e0201939.

48. Dinges DF, Powell JW. 1985. Microcomputer analyses of performance on a portable, simple visual RT task during sustained operations. Behavior Research Methods, Instruments, & Computers 17:652–655.

49. Basner M, Mollicone D, Dinges DF. 2011. Validity and sensitivity of a brief psychomotor vigilance test (PVT-B) to total and partial sleep deprivation. Acta Astronautica 69:949–959.

50. Anderson KA, Points III GL, Donald CE, Dixon HM, Scott RP, Wilson G, Tidwell LG, Hoffman PD, Herbstman JB, O’Connell SG. 2017. Preparation and performance features of wristband samplers and considerations for chemical exposure assessment. Journal of Exposure Science & Environmental Epidemiology 27:551–559.

51. Hammel SC, Hoffman K, Webster TF, Anderson KA, Stapleton HM. 2019. Discovery of common chemical exposures across three continents using silicone wristbands. Royal Society Open Science 6:181836.

52. Dixon HM, Bramer LM, Scott RP, Calero L, Holmes D, Gibson EA, Cavalier HM, Rohlman D, Miller RL, Calafat AM, Kincl L, Waters KM, Herbstman JB, Anderson KA. 2022. Evaluating predictive relationships between wristbands and urine for assessment of personal PAH exposure. Environment International 163:107226.

53. Kundu P, Torres ERS, Stagaman K, Kasschau K, Okhovat M, Holden S, Ward S, Nevonen KA, Davis BA, Saito T, Saido TC, Carbone L, Sharpton TJ, Raber J. 2021. Integrated analysis of behavioral, epigenetic, and gut microbiome analyses in App NL-G-F, App NL-F, and wild type mice. Scientific Reports 11.

54. Caporaso JG, Lauber CL, Walters WA, Berg-Lyons D, Huntley J, Fierer N, Owens SM, Betley J, Fraser L, Bauer M, Gormley N, Gilbert JA, Smith G, Knight R. 2012. Ultra-high-throughput microbial community analysis on the Illumina HiSeq and MiSeq platforms. ISME J 6:1621–1624.

55. Callahan BJ, McMurdie PJ, Rosen MJ, Han AW, Johnson AJA, Holmes SP. 2016. DADA2: High-resolution sample inference from Illumina amplicon data. Nat Methods 13:581–583.

56. Hammer AJ, Gaulke CA, Garcia-Jaramillo M, Leong C, Morre J, Sieler MJ, Stevens JF, Jiang Y, Maier CS, Kent ML, Sharpton TJ. 2024. Gut microbiota metabolically mediate intestinal helminth infection in zebrafish. mSystems 9:e0054524.

57. Dixon P. 2003. VEGAN, a package of R functions for community ecology. Journal of Vegetation Science 14:927–930.

58. Mallick H, Rahnavard A, McIver LJ, Ma S, Zhang Y, Nguyen LH, Tickle TL, Weingart G, Ren B, Schwager EH, Chatterjee S, Thompson KN, Wilkinson JE, Subramanian A, Lu Y, Waldron L, Paulson JN, Franzosa EA, Bravo HC, Huttenhower C. 2021. Multivariable association discovery in population-scale meta-omics studies. PLOS Computational Biology 17:e1009442.

59. Wickham H. 2016. ggplot2: Elegant Graphics for Data Analysis. Springer.

60. Hou J, Sun H, Zhou Y, Zhang Y, Yin W, Xu T, Cheng J, Chen W, Yuan J. 2017. Impacts of low socioeconomic status and polycyclic aromatic hydrocarbons exposure on lung function among a community-based Chinese population. Science of The Total Environment 574:1095–1103.

61. Wang L, Zhang L, Hou J, Liu C, Zhang Y, Zhou Y, Yin W, Li S, Yuan J. 2019. Mediating factors explaining the associations between polycyclic aromatic hydrocarbons exposure, low socioeconomic status and diabetes: A structural equation modeling approach. Science of The Total Environment 648:1476–1483.

62. Jongeneelen FJ. 2001. Monitoring human occupational and environmental exposures to polycyclic aromatic compounds. Annals of Work Exposures and Health 47:349–378.

63. Alomirah H, Al-Zenki S, Zaghloul S, Sawaya W, Ahmed N. 2011. Concentrations and dietary exposure to polycyclic aromatic hydrocarbons (PAHs) from grilled and smoked foods. Food Control 22:2028–2035.

64. Berge G, Mollerup S, Övrebo S, Hewer A, Phillips DH, Eilertsen E, Haugen A. 2004. Role of estrogen receptor in regulation of polycyclic aromatic hydrocarbon metabolic activation in lung. Lung Cancer 45:289–297.

65. Guo H, Huang K, Zhang X, Zhang W, Guan L, Kuang D, Deng Q, Deng H, Zhang X, He M, Christiani DC, Wu T. 2014. Women are more susceptible than men to oxidative stress and chromosome damage caused by polycyclic aromatic hydrocarbons exposure. Environmental and Molecular Mutagenesis 55:472–481.

66. Cross CP, Copping LT, Campbell A. 2011. Sex differences in impulsivity: A meta-analysis. Psychological Bulletin 137:97–130.

67. Weafer J, de Wit H. 2014. Sex differences in impulsive action and impulsive choice. Addictive Behaviors 39:1573–1579.

68. Weinstein A, Dannon P. 2015. Is impulsivity a male trait rather than female trait? Exploring the sex difference in impulsivity. Current Behavioral Neuroscience Reports 2:9–14.

69. Arteaga-Henríquez G, Simon MS, Burger B, Weidinger E, Wijkhuijs A, Arolt V, Birkenhäger TK, Musil R, Müller N, Drexhage HA. 2020. Treating impulsivity with probiotics in adults (PROBIA): study protocol of a multicenter, double-blind, randomized, placebo-controlled trial. Trials 21:161.

70. Frankot MA, O’Hearn CM, Blancke AM, Rodriguez B, Pechacek KM, Gandhi J, Hu G, Martens KM, Vonder Haar C. 2023. Acute gut microbiome changes after traumatic brain injury are associated with chronic deficits in decision-making and impulsivity in male rats. Behavioral Neuroscience 137:15–28.

71. Liu W, Zhang X, Zhou B, Shang L, Zhang Y, Liang L, Yang Z, Liu X, Liu T, Yuan Y, Zhang X, Liang S. 2024. Associations between impulsivity and fecal microbiota in individuals abstaining from methamphetamine. CNS Neuroscience & Therapeutics 30:e14580.

72. Langmajerová M, Roubalová R, Šebela A, Vevera J. 2023. The effect of microbiome composition on impulsive and violent behavior: A systematic review. Behavioural Brain Research 440:114266.

73. Mollerup S, Berge G, Baera R, Skaug V, Hewer A, Phillips DH, Stangeland L, Haugen A. 2006. Sex differences in risk of lung cancer: Expression of genes in the PAH bioactivation pathway in relation to smoking and bulky DNA adducts. International Journal of Cancer 119:741–744.

74. Uppstad H, Osnes GH, Cole KJ, Phillips DH, Haugen A, Mollerup S. 2011. Sex differences in susceptibility to PAHs is an intrinsic property of human lung adenocarcinoma cells. Lung Cancer 71:264–270.

75. Vondráček J, Pivnička J, Machala M. 2018. Polycyclic aromatic hydrocarbons and disruption of steroid signaling. Current Opinion in Toxicology 11–12:27–34.

76. Fernandes D, Porte C. 2013. Hydroxylated PAHs alter the synthesis of androgens and estrogens in subcellular fractions of carp gonads. Science of The Total Environment 447:152–159.

77. Yang Q, Zhao Y, Qiu X, Zhang C, Li R, Qiao J. 2015. Association of serum levels of typical organic pollutants with polycystic ovary syndrome (PCOS): a case–control study. Human Reproduction 30:1964–1973.

78. Feng Q, Mu G, Zhang X, Wang H, Zhou L, Cao L, Chen W, Yuan J. 2023. Focusing on testosterone levels in male: A half-longitudinal study of polycyclic aromatic hydrocarbon exposure and diastolic blood pressure in coke oven workers. Environmental Pollution 329:121614.

79. Shields GS, Ivory SL, Telzer EH. 2019. Three-month cumulative exposure to testosterone and cortisol predicts distinct effects on response inhibition and risky decision-making in adolescents. Psychoneuroendocrinology 110:104412.

80. Zhuang J-Y, Wang J-X, Lei Q, Zhang W, Fan M. 2020. Neural basis of increased cognitive control of impulsivity during the mid-luteal phase relative to the late follicular phase of the menstrual cycle. Frontiers in Human Neuroscience 14:568399.

81. Gui X, Yang Z, Li MD. 2021. Effect of cigarette smoke on gut microbiota: State of knowledge. Frontiers in Physiology 12.

82. Engen PA, Green SJ, Voigt RM, Forsyth CB, Keshavarzian A. 2015. The gastrointestinal microbiome: Alcohol effects on the composition of intestinal microbiota. Alcohol Research: Current Reviews 37:223–236.

83. Miclotte L, Van de Wiele T. 2020. Food processing, gut microbiota and the globesity problem. Critical Reviews in Food Science and Nutrition 60:1769–1782.

84. Qiu F, Liang CL, Liu H, Zeng YQ, Hou S, Huang S, Lai X, Dai Z. 2017. Impacts of cigarette smoking on immune responsiveness: Up and down or upside down? Oncotarget 8:268–284.

85. Sarkar D, Jung MK, Wang HJ. 2015. Alcohol and the immune system. Alcohol Research: Current Reviews 37:153–155.

86. Goldman R, Enewold L, Pellizzari E, Beach JB, Bowman ED, Krishnan SS, Shields PG. 2001. Smoking increases carcinogenic polycyclic aromatic hydrocarbons in human lung tissue. Cancer Research 61:6367–6371.

87. Martin AM, Young RL, Leong L, Rogers GB, Spencer NJ, Jessup CF, Keating DJ. 2019. The gut microbiome regulates host glucose homeostasis via peripheral serotonin. Proceedings of the National Academy of Sciences 116:19802–19804.

88. Yano JM, Yu K, Donaldson GP, Shastri GG, Ann P, Ma L, Nagler CR, Ismagilov RF, Mazmanian SK, Hsiao EY. 2015. Indigenous bacteria from the gut microbiota regulate host serotonin biosynthesis. Cell 161:264–276.

89. Reigstad CS, Salmonson CE, Rainey III JF, Szurszewski JH, Linden DR, Sonnenburg JL, Farrugia G, Kashyap PC. 2015. Gut microbes promote colonic serotonin production through an effect of short-chain fatty acids on enterochromaffin cells. FASEB Journal 29:1395–1403.

90. O’Mahony SM, Clarke G, Borre YE, Dinan TG, Cryan JF. 2015. Serotonin, tryptophan metabolism and the brain-gut-microbiome axis. Behavioural Brain Research 277:32–48.

91. Marazziti D, Baroni S, Masala I, Golia F, Consoli G, Massimetti G, Picchetti M, Catena Dell’osso M, Giannaccini G, Betti L, Lucacchini A, Ciapparelli A. 2010. Impulsivity, gender, and the platelet serotonin transporter in healthy subjects. Neuropsychiatric Disease and Treatment 6:9–15.

92. De Filippo C, Cavalieri D, Di Paola M, Ramazzotti M, Poullet JB, Massart S, Collini S, Pieraccini G, Lionetti P. 2010. Impact of diet in shaping gut microbiota revealed by a comparative study in children from Europe and rural Africa. Proc Natl Acad Sci U S A 107:14691–14696.

93. Wu GD, Chen J, Hoffmann C, Bittinger K, Chen Y-Y, Keilbaugh SA, Bewtra M, Knights D, Walters WA, Knight R, Sinha R, Gilroy E, Gupta K, Baldassano R, Nessel L, Li H, Bushman FD, Lewis JD. 2011. Linking Long-Term Dietary Patterns with Gut Microbial Enterotypes. Science 334:105–108.

94. Bravo JA, Forsythe P, Chew MV, Escaravage E, Savignac HM, Dinan TG, Bienenstock J, Cryan JF. 2011. Ingestion of Lactobacillus strain regulates emotional behavior and central GABA receptor expression in a mouse via the vagus nerve. Proc Natl Acad Sci U S A 108:16050–16055.

95. Wexler HM. 2007. Bacteroides: the good, the bad, and the nitty-gritty. Clin Microbiol Rev 20:593–621.

96. Rouskas K, Mamalaki E, Ntanasi E, Pantoura M, Anezaki M, Emmanouil C, Novau-Ferré N, Bulló M, Dimas AS, Papandreou C, Yannakoulia M, Argiriou A, Scarmeas N. 2025. Gut Microbiome Alterations in Mild Cognitive Impairment: Findings from the ALBION Greek Cohort. Microorganisms 13:2112.

97. Lin S-H, Lin R-J, Chu C-L, Chen Y-L, Fu S-C. 2025. Associations Between Gut Microbiota Composition and Impulse Control Disorders in Parkinson’s Disease. Int J Mol Sci 26:6146.

98. Shoubridge AP, Carpenter L, Flynn E, Papanicolas LE, Collins J, Gordon D, Lynn DJ, Whitehead C, Leong LEX, Cations M, De Souza DP, Narayana VK, Choo JM, Wesselingh SL, Crotty M, Inacio MC, Ivey K, Taylor SL, Rogers GB. 2025. Severe Cognitive Decline in Long-term Care Is Related to Gut Microbiome Production of Metabolites Involved in Neurotransmission, Immunomodulation, and Autophagy. J Gerontol A Biol Sci Med Sci 80:glaf053.

99. Jangi S, Gandhi R, Cox LM, Li N, von Glehn F, Yan R, Patel B, Mazzola MA, Liu S, Glanz BL, Cook S, Tankou S, Stuart F, Melo K, Nejad P, Smith K, Topçuolu BD, Holden J, Kivisäkk P, Chitnis T, De Jager PL, Quintana FJ, Gerber GK, Bry L, Weiner HL. 2016. Alterations of the human gut microbiome in multiple sclerosis. Nature Communications 7:12015.

100. Dunalska A, Saramak K, Szejko N. 2023. The Role of Gut Microbiome in the Pathogenesis of Multiple Sclerosis and Related Disorders. Cells 12:1760.

